# Alcohol, Tobacco, and HPV exposure synergistically regulate promoter methylation of PD-L1 related genes in Head and Neck Squamous Cell Carcinoma

**DOI:** 10.64898/2026.05.26.728017

**Authors:** Omar Mokhashi, Lan Gao, Ruomin Xin, Riya Chhabra, Stella Hale, Jessica Wang-Rodriguez, Weg M. Ongkeko

## Abstract

Although alcohol, tobacco and human papillomavirus (HPV) are established risk factors for head and neck squamous cell carcinoma (HNSCC), the molecular effects of their co-occurrence remain poorly understood. DNA methylation may link exposure history to regulation of the PD-1/PD-L1 immune checkpoint pathway, whose therapeutic blockade benefits only a subset of patients with recurrent or metastatic HNSCC. We stratified primary tumors from The Cancer Genome Atlas HNSCC cohort by all combinations of these exposures and examined promoter methylation across seven interferon-γ–PD-L1 signaling genes at single-CpG resolution, with region-level validation. Triple-exposure tumors exhibited coordinated methylation changes involving CD274, IRF1, JAK1, PDCD1, MIR33A and MIR34A, with differentially methylated regions overlapping PDCD1, MIR34AHG and JAK1. CD274 promoter methylation is inversely correlated with gene expression. Significant methylation differences occurred only in HPV-positive subgroups, suggesting HPV may be necessary for this phenotype, while concurrent exposures may modify its extent. Transcriptomic analysis and immune deconvolution further suggested greater estimated immune infiltration alongside downregulation of innate and adaptive immune activation pathways, consistent with an inflamed but functionally suppressed tumor microenvironment. These findings suggest combined exposure history may help contextualize checkpoint pathway regulation in HNSCC.

## Introduction

Head and neck squamous cell carcinoma is among the most common malignancies worldwide, and the majority of cases are attributable to three exposures: chronic alcohol consumption, tobacco use, and infection with high-risk human papillomavirus (HPV) [1–3]. Although each of these is a well-established risk factor, they are conventionally treated as alternative etiologies, and HNSCC is routinely dichotomized into HPV-positive and HPV-negative disease for prognostic and therapeutic purposes [4,5]. In practice, however, these exposures frequently co-occur within the same patient, and a substantial proportion of HPV-positive tumors arise in individuals who also drink and smoke [4,6]. The molecular consequences of this overlap have received comparatively little attention.

Subsequently, while immune checkpoint inhibitors targeting PD-1 have become standard therapy in recurrent and metastatic HNSCC, objective response rates remain below twenty per cent [7,8]. PD-L1 combined positive score is the principal approved biomarker, yet it stratifies response imperfectly, and many patients whose tumors express PD-L1 derive no benefit [7,8]. This has prompted increasing interest in the regulatory machinery upstream of PD-L1 rather than in PD-L1 abundance alone.

The dominant physiological route to tumor PD-L1 expression is adaptive and well characterized [9]. Interferon-γ released by infiltrating lymphocytes engages IFNGR, activating JAK1 and JAK2, which in turn phosphorylate STAT1 and induce IRF1, the transcription factor that transactivates CD274 [10]. Disruption anywhere along this axis uncouples PD-L1 expression from immune pressure, and loss-of-function mutations in JAK1 and JAK2 are established mechanisms of acquired resistance to checkpoint blockade in melanoma and mismatch repair-deficient tumors [11,12]. Whether this axis can be disrupted through non-genetic means in HNSCC, and whether exposure history contributes to such disruption, is not yet definitively known.

DNA methylation represents a plausible mediator. Smoking and alcohol each leave reproducible methylation signatures in human tissue [13,14], HPV-positive HNSCC carries a globally distinct and broadly hypermethylated genome [5,15], and promoter methylation of CD274 modulates PD-L1 expression in several malignancies [16,17]. Whether combinations of exposure shape checkpoint pathway output epigenetically have not, however, been examined systematically.

Checkpoint output is also shaped outside the interferon-γ cascade. PD-L1 signals through PD-1, encoded by PDCD1, whose promoter methylation has been associated with outcome in HNSCC [18], and PD-L1 is regulated post-transcriptionally by miR-34a, which directly targets the CD274 transcript [19], and miR-33a, which has been linked to PD-L1 status in lung adenocarcinoma [20]; MIR34A is also a frequent target of promoter hypermethylation in cancer [21]. We therefore examined a panel comprising JAK1, JAK2 and IRF1, CD274 and PDCD1, and MIR33A and MIR34A.

In this study, we stratified 498 primary tumors from the TCGA HNSCC cohort into subgroups reflecting all possible combinations of alcohol, tobacco and HPV exposure, and we characterized promoter methylation across this candidate panel at both single-CpG and region level. We then related methylation to transcript abundance in matched samples, and placed these findings in the context of the transcriptional and immune-microenvironmental landscape of each exposure subgroup.

## Materials and Methods

### Data acquisition and cohort definition

Multi-omic data for the TCGA-HNSC cohort [5] were accessed through the Genomic Data Commons using the TCGAbiolinks R package [22]. DNA methylation data were retrieved as GDC-harmonized Level-3 beta values from the Illumina Infinium HumanMethylation450 BeadChip (hg38), together with matched clinical metadata and mRNA and miRNA sequencing reads. Only primary tumor samples were retained. Methylation barcodes were matched to clinical metadata for all 498 patients, and a matched mRNA raw count matrix of 59,900 genes across 487 samples was retained for the methylation–expression analysis.

### Exposure classification

Each patient was assigned to one of eight subgroups reflecting all combinations of alcohol exposure (TCGA clinical alcohol history field), tobacco exposure (any recorded smoking history) and HPV status (TCGA clinical annotation): Group A, no exposure (n = 82); B, alcohol only (n = 91); C, tobacco only (n = 65); D, HPV only (n = 13); E, alcohol and tobacco (n = 174); F, alcohol and HPV (n = 28); G, tobacco and HPV (n = 0); and H, all three exposures (n = 45). Group G was empty and excluded from all comparisons (Fig.1b).

### DNA methylation processing and probe filtering

Of the 485,577 probes represented on the array, any probe with a missing value in one or more samples was removed, leaving 334,609 probes with complete data. Probes affected by an underlying single-nucleotide polymorphism, probes with known cross-reactivity [23], and probes mapping to the sex chromosomes were then removed using the rmSNPandCH function in DMRcate [24], leaving 325,410 probes for analysis. A targeted candidate panel of 51 probes was defined prior to analysis across seven genes of the interferon-γ–PD-L1 signaling axis: *CD274*, *PDCD1*, *IRF1*, *JAK1*, *JAK2*, *MIR33A* and *MIR34A*. Probes were selected by Illumina UCSC RefGene annotation group, comprising those annotated to the TSS1500 (n = 17), TSS200 (n = 17), 5′UTR (n = 13) or first exon (n = 4) regions of these genes. Probe representation per gene was uneven, reflecting array coverage: *JAK1* (n = 19), *MIR33A* and *PDCD1* (n = 8 each), *IRF1* (n = 7), and *CD274*, *JAK2* and *MIR34A* (n = 3 each). Throughout this work, "promoter methylation" refers to methylation at probes within these annotation groups. Beta values were converted to M-values as M = log₂[(β + 0.001)/(1 − β + 0.001)] [25]. M-values were used for all statistical testing, while beta values were retained for visualization.

### Differential methylation analysis

Six contrasts, each comparing one exposure group with unexposed controls, were tested using a ∼0 + exposure_group design in limma [26] (lmFit, eBayes, topTable). Benjamini–Hochberg correction [27] was applied both genome-wide across all 325,410 probes, as well as separately within the 51 candidate probes; main figures report the targeted correction, and genome-wide results are provided as supplementary material.

### Region-level confirmation

For the triple-exposure versus control contrast, differentially methylated regions were called genome-wide with DMRcate [24] (cpg.annotate; dmrcate, λ = 1000 bp; extractRanges; hg38) and intersected with the candidate gene list. Because regions were not restricted to promoter intervals, a region assigned to a candidate gene may fall outside its promoter probes; promoter-level and region-level results therefore refer to distinct intervals and are reported separately.

### Methylation–expression coupling

The mRNA count matrix was collapsed by gene symbol; 473 symbols were duplicated, none of which corresponded to a candidate gene. Counts were normalized to log₂(CPM + 1). Spearman rank correlation was computed between each candidate gene’s promoter probes and the corresponding gene’s expression across the 487 matched patients, with Benjamini–Hochberg correction applied within each gene. *MIR33A* and *MIR34A* were not included in this analysis, as mature microRNAs are not represented in the mRNA-seq count matrix.

### RNA sequencing and differential expression

Raw RNA-sequencing reads underwent quality control assessment using FastQC, with adapter trimming performed using Cutadapt [28]. Trimmed reads were aligned to the human reference genome (GRCh38) using the STAR aligner [29], and gene-level read counts were obtained using featureCounts [30] for mRNA and miRDeep2 [31] for miRNA. These processing steps were executed via SLURM batch jobs submitted to the SDSC Expanse supercomputer. Differential expression analysis was performed using DESeq2 [32], with each exposure subgroup compared independently against the unexposed control group (Group A). Genes were considered significantly differentially expressed at an adjusted p value (padj) below 0.05, with p values corrected for multiple comparisons using the Benjamini–Hochberg procedure [27]. Hierarchical clustering of normalized expression values was performed to visualize transcriptomic patterns across exposure subgroups.

### Gene set enrichment analysis

Genes were ranked by their differential expression statistics from the triple-exposure versus control comparison and tested against Gene Ontology Biological Process gene sets by gene set enrichment analysis [33] using the clusterProfiler and enrichplot packages [34]. Enrichment significance was assessed by permutation, with Benjamini–Hochberg adjustment of enrichment p values.

### Immune deconvolution

Immune populations were estimated from bulk RNA-seq using TIMER2.0 [35] (http://timer.cistrome.org/): CD8⁺ T cells, activated memory CD4⁺ T cells and regulatory T cells with CIBERSORT-ABS [36], M1 and M2 macrophages with xCell [37], and natural killer cells with quanTIseq [38]. Each exposure group was compared with controls by two-sided Wilcoxon rank-sum test, with Benjamini–Hochberg correction across populations.

### Somatic mutation and CNV analysis

Somatic mutation data from the GDC were compared between triple-exposure and control tumors for genes of the interferon-γ–PD-L1 axis using maftools [39] and visualized with coOncoplot. Given the low mutation frequencies at these loci, this analysis was treated as a test of whether somatic alteration offers an alternative explanation for the pathway-level differences rather than as a discovery analysis. Discrete gene-level copy number calls for TCGA-HNSC were obtained from cBioPortal [40] and categorized as deletion, diploid, gain or amplification. Gene-wise mutation and copy number frequencies were compared using Fisher’s exact test with Benjamini–Hochberg correction, and odds ratios with 95% confidence intervals were calculated for mutation frequencies.

### Statistical analysis and software

All analyses were performed in R version 4.5.2. Statistical significance was defined as an adjusted p value below 0.05 throughout, unless otherwise specified. Package versions are listed in the supplementary material.

## Results

### Cohort composition and exposure distribution

Of the 498 primary HNSCC tumors retained after quality control, all were successfully assigned to an exposure category on the basis of clinical alcohol history, smoking history and HPV status (Fig.1A). Alcohol and tobacco exposure co-occurred frequently, with combined alcohol and tobacco exposure representing the largest subgroup in the cohort (n = 174). HPV-positive tumors were comparatively uncommon and were almost always accompanied by at least one behavioral exposure: of the 86 HPV-positive tumors, only fi13 carried HPV exposure alone, while 28 carried HPV with alcohol and 45 carried all three exposures (Fig.1B). Notably, no patient in this cohort was classified as tobacco-exposed and HPV-positive in the absence of alcohol exposure, and Group G was therefore empty and excluded from all subsequent comparisons. This distribution illustrates that the conventional dichotomy between HPV-driven and behaviorally driven HNSCC does not reflect how these exposures are distributed in practice, and it defines the seven populated subgroups used throughout the remainder of this study.

**Figure 1.**
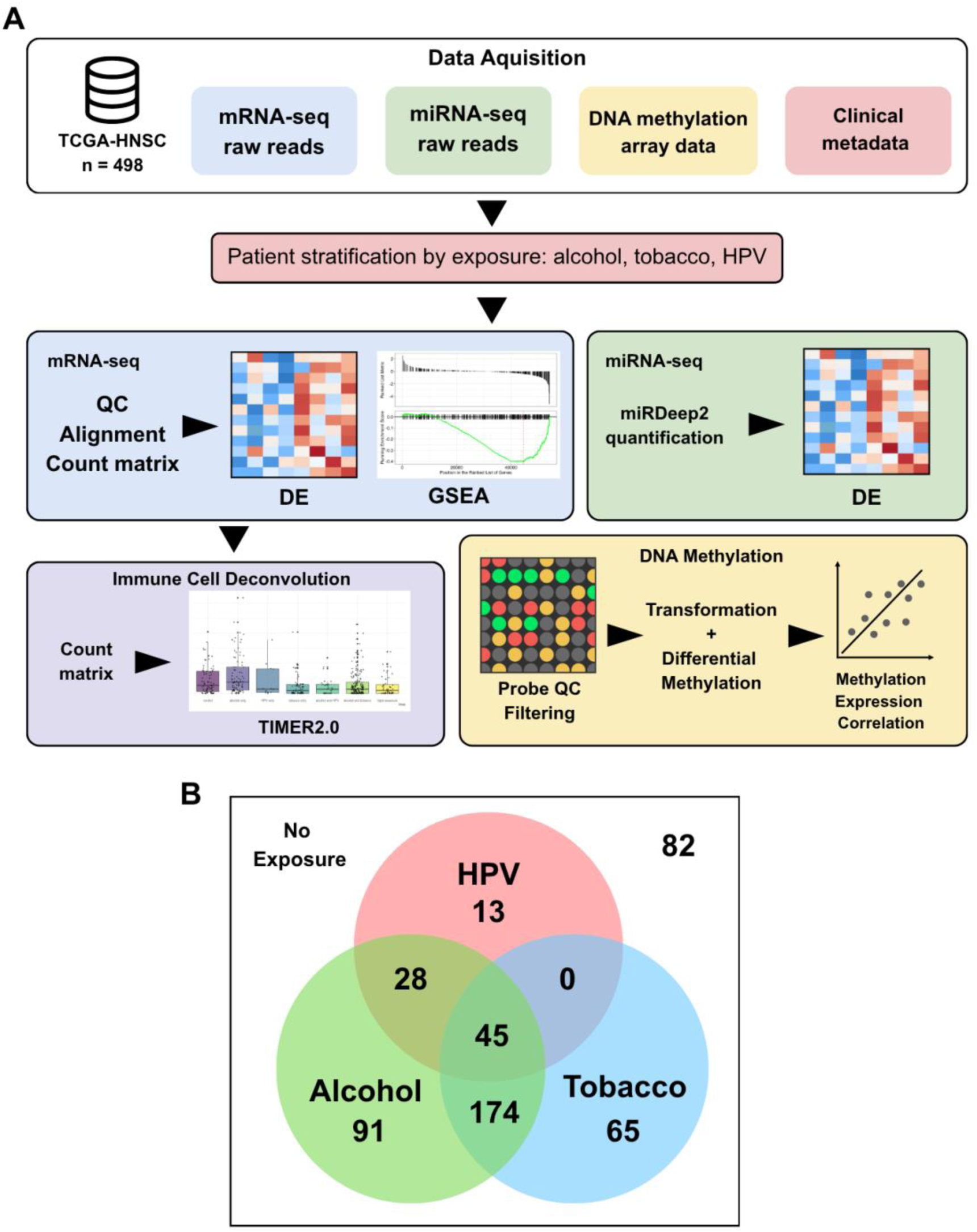
Study design and cohort composition. (A) Analytical workflow. TCGA-HNSC mRNA and miRNA sequencing reads, DNA methylation array data and clinical metadata were retrieved, and patients were stratified by alcohol, tobacco and HPV exposure before differential expression, gene set enrichment, immune deconvolution and DNA methylation analyses. (B) Distribution of the 498 primary tumors across all exposure combinations.

### Promoter methylation of the checkpoint axis is restricted to HPV-containing exposure groups

To determine whether exposure combinations are associated with methylation of the interferon-γ–PD-L1 signaling axis, we examined 51 promoter probes across seven candidate genes in each exposure group relative to unexposed controls (Fig. 2). Differential methylation was concentrated in the triple-exposure group, in which twelve probes reached significance. *CD274* was hypermethylated at cg19724470 (difference in M-value, ΔM = +0.63; FDR = 0.0039), while *PDCD1* and *MIR33A* were hypomethylated at cg20805133 (ΔM = −0.85, FDR = 0.0039) and cg24260710 (ΔM = −0.59, FDR = 1.5 × 10⁻⁶), respectively (Figs 2 and 4A). *MIR34A* was hypermethylated at cg00909706 (ΔM = +0.24, FDR = 0.025; Figs 2 and 4A), and smaller but significant hypermethylation was observed at *JAK1* (cg14425609, ΔM = +0.15, FDR = 0.012) and *IRF1* (cg00578828, ΔM = +0.16, FDR = 0.035) (Fig. 2). *JAK2* did not reach significance in any group, approaching the threshold only under triple exposure (cg03508284, FDR = 0.079; Fig. 2). Six of the twelve significant probes were at *PDCD1*, indicating that change at this locus was distributed across neighboring sites.

**Figure 2.**
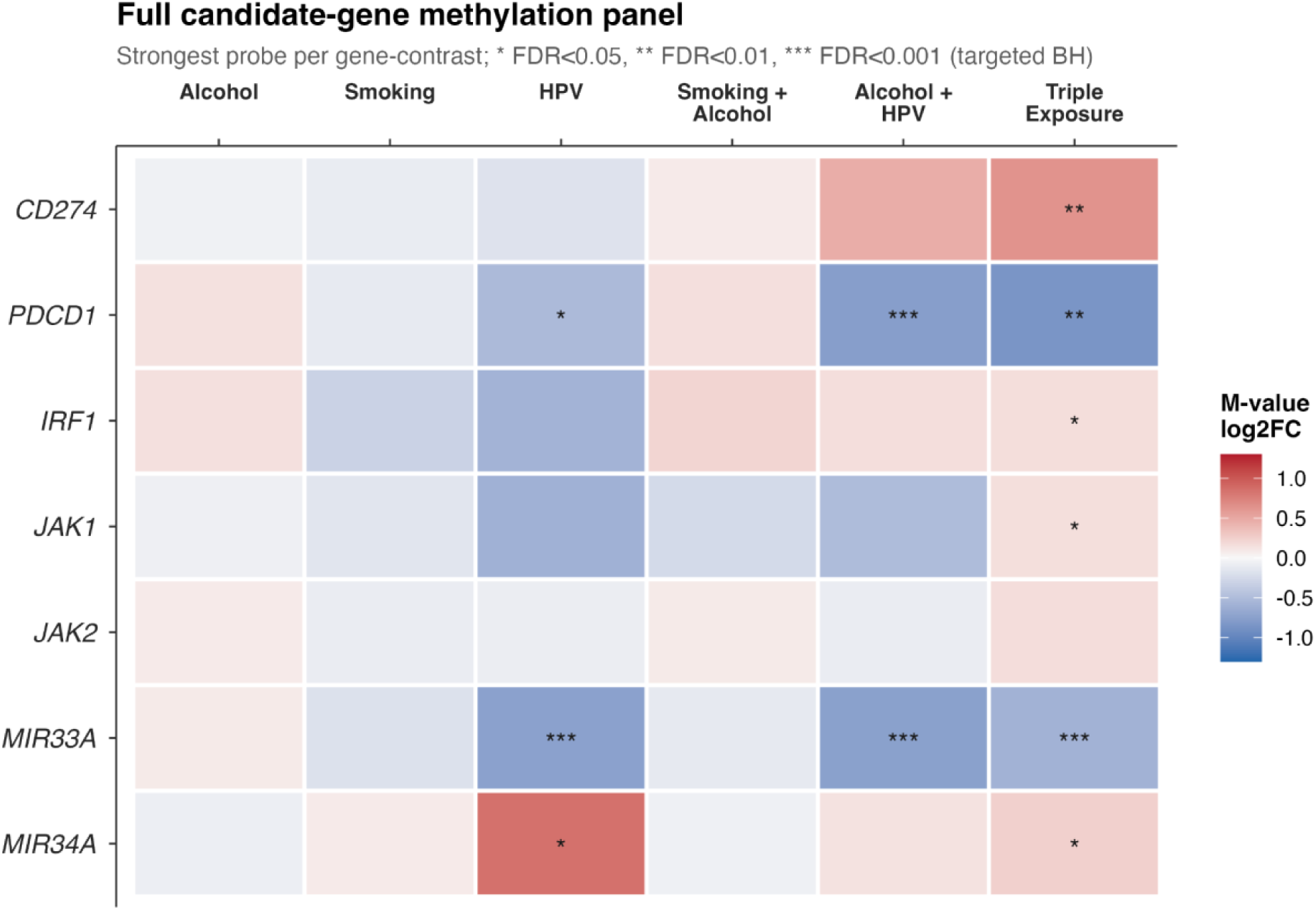
Promoter methylation across the full interferon-γ–PD-L1 candidate panel. Differential promoter methylation for all seven candidate genes across six exposure contrasts, each compared with the unexposed control group. Color indicates the M-value log₂ fold change, with red denoting hypermethylation and blue hypomethylation relative to control. The strongest probe per gene–contrast pair is shown. Asterisks denote Benjamini–Hochberg adjusted significance within the 51-probe candidate panel (*FDR < 0.05, **FDR < 0.01, ***FDR < 0.001). No probe reached significance in the alcohol-only, smoking-only or combined smoking and alcohol contrasts, and *JAK2* showed no significant change in any contrast.

A comparable pattern was observed in the two other subgroups containing HPV (Fig. 2). Alcohol and HPV co-exposure was associated with hypomethylation at *MIR33A* (cg24260710, ΔM = −0.74, FDR = 1.4 × 10⁻⁷) and *PDCD1* (cg17322655, ΔM = −0.76, FDR = 2.3 × 10⁻⁵) (Figs 2 and 4A), with ten probes reaching significance in total and the largest single effect in the panel observed at *PDCD1* cg20805133 (ΔM = −1.21). HPV exposure alone was associated with three significant probes despite the small size of this subgroup (n = 13): hypomethylation at *MIR33A* (ΔM = −0.74, FDR = 0.00066) and *PDCD1* (cg18096388, ΔM = −0.52, FDR = 0.012), and hypermethylation at *MIR34A* (cg24501230, ΔM = +0.85, FDR = 0.012) (Figs 2 and 4A). The largest *MIR34A* effect was therefore observed in the HPV-only group, compared with ΔM = +0.24 at cg00909706 under triple exposure (Figs 2 and 4A).

No significant differential methylation was detected at any of the 51 candidate probes in the alcohol-only, tobacco-only or combined alcohol and tobacco subgroups (Figs 2 and 4A), despite the combined alcohol and tobacco group being the largest in the cohort (n = 174). None of these contrasts approached significance; the smallest FDR values were 0.77, 0.37 and 0.61, respectively.

The same restriction was evident genome wide. Of the 325,410 retained probes, the number reaching genome-wide significance was 2 in the alcohol-only contrast, 14 in the tobacco-only contrast and 64 in the combined alcohol and tobacco contrast, compared with 23,760 in the HPV-only contrast, 44,480 under alcohol and HPV co-exposure, and 62,026 under triple exposure. The three HPV-negative exposure groups therefore produced almost no detectable methylation change at any locus, including the largest subgroup in the cohort, while every HPV-containing group produced tens of thousands of altered probes. The number of altered probes was greatest under triple exposure, although these counts also scale with subgroup size (n = 13, 28 and 45). Across the candidate panel, triple exposure was likewise the only contrast in which six of the seven candidate genes reached significance (Fig. 2).

### Region-level analysis supports methylation changes at three candidate loci

To test whether probe-level findings reflected coordinated change across neighboring CpGs, we intersected genome-wide differentially methylated regions from the triple-exposure contrast with the candidate genes (Fig. 3). Three were represented: *MIR34AHG*, the host gene of *MIR34A* (two CpGs, mean Δβ = +0.115; chr1:9,154,250–9,154,254; HMFDR = 1.5 × 10⁻⁵), *PDCD1* (three CpGs, mean Δβ = +0.111; chr2:241,851,343–241,851,501; HMFDR = 2.2 × 10⁻⁵) and *JAK1* (ten CpGs, mean Δβ = +0.026; chr1:64,935,606–64,937,528; HMFDR = 0.0078) (Fig. 3).

**Figure 3.**
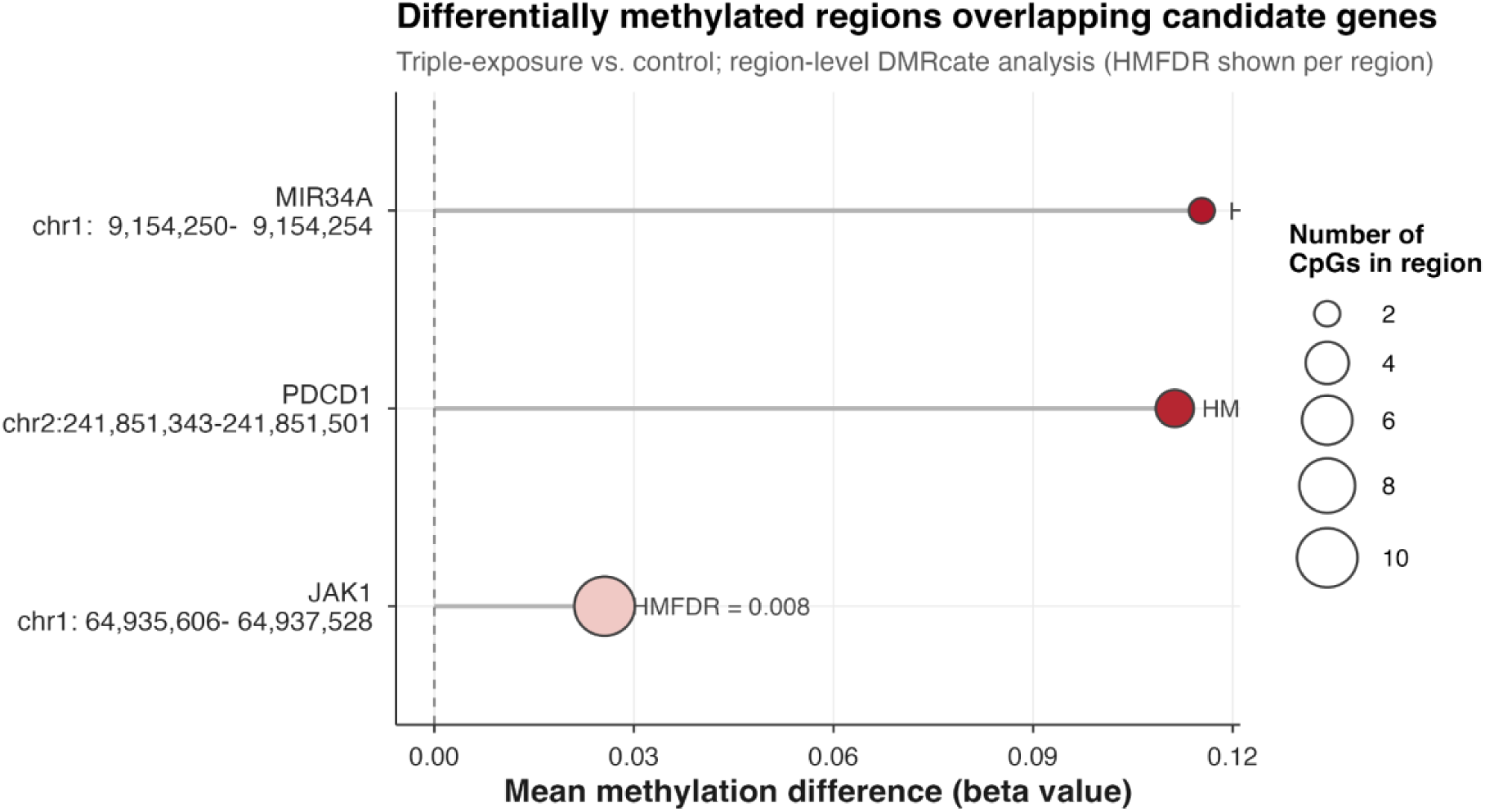
Differentially methylated regions overlapping candidate genes in triple-exposure tumors. Region-level analysis of the triple-exposure versus control contrast, performed with DMRcate across all retained probes and intersected with the candidate gene list. The x-axis shows the mean difference in beta value between groups, with positive values indicating higher methylation in triple-exposure tumors. Point size indicates the number of CpG sites contributing to each region. Region coordinates are given on genome build hg38, and the harmonic mean false discovery rate (HMFDR) is shown for each region.

The *JAK1* region spanned the widest interval (1,923 bp) but had the smallest effect (Fig. 3). *CD274* was not represented among the significant regions (Fig. 3), and its promoter hypermethylation is therefore supported at the probe level only (Figs 2 and 4A). The *PDCD1* region identified here is hypermethylated relative to control (Fig. 3), whereas the promoter probes examined at the probe level were hypomethylated (Figs 2 and 4A). Because regions were not restricted to promoter intervals, these results refer to distinct intervals and are reported separately [41].

**Figure 4.**
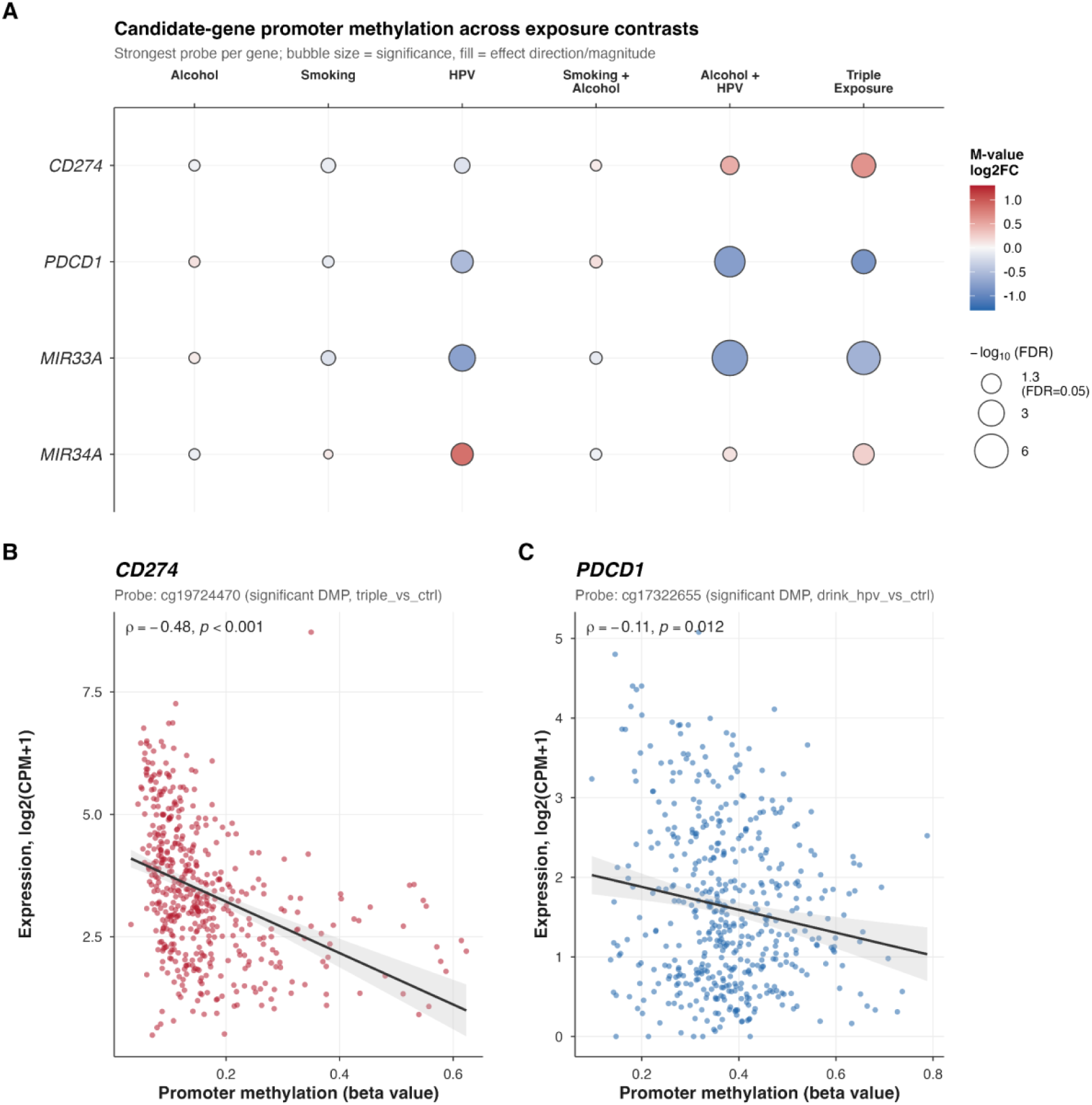
Methylation of checkpoint genes and its relationship to transcript abundance. (**A**)Differential promoter methylation for the four candidate genes with the largest significant changes across exposure contrasts; fill color encodes effect direction and magnitude, point size encodes significance (−log₁₀ FDR). These contrasts are a subset of Figure 2, shown here to orient panels B and C. (**B**) Spearman correlation between methylation at cg19724470 and *CD274* expression across 487 matched patients (ρ = −0.48, p < 2.2 × 10⁻¹⁶); this was the most significantly hypermethylated *CD274* probe in the triple-exposure contrast. (**C**) The same for cg17322655 and *PDCD1* expression (ρ = −0.11, adjusted p = 0.025), the most significantly differentially methylated *PDCD1* probe in the alcohol and HPV contrast. This probe was the most significantly differentially methylated *PDCD1* probe in the alcohol and HPV versus control contrast. Expression is shown as log₂(CPM + 1); methylation is shown as beta value. Shaded bands indicate 95% confidence intervals around the fitted line.

### Promoter methylation at CD274 is inversely related to CD274 expression

Given that the functional consequence of promoter methylation depends on its effect on transcription, we related methylation at each candidate promoter probe to transcript abundance across the 487 patients with matched expression data, covering 40 probes across the five candidate genes represented in the mRNA-seq matrix.

Methylation at cg19724470, the most significantly hypermethylated *CD274* probe in triple-exposure tumors (Figs 2 and 4A), was inversely correlated with *CD274* expression (Spearman ρ = −0.48, padj < 0.001; Fig. 4B), the strongest methylation–expression relationship in the panel. Two further *CD274* probes showed weaker inverse relationships (cg14305799, ρ = −0.27; cg02823866, ρ = −0.16; both padj < 0.01), indicating that the relationship is not confined to a single site. Together these provide a plausible transcriptional route by which the observed methylation changes may influence checkpoint pathway output.

The corresponding relationship for *PDCD1* at cg17322655 was considerably weaker (ρ = −0.11, padj = 0.025; Fig. 4C), consistent with *PDCD1* being expressed predominantly by infiltrating lymphocytes rather than tumor cells, so that bulk methylation and transcript abundance derive only partly from the same cells.

The probes at which *IRF1* and *JAK1* reached significance (Fig. 2) showed no relationship with their own expression (cg00578828, ρ = +0.03, padj = 0.55; cg14425609, ρ = +0.00, padj = 0.95), and we therefore do not interpret these loci as carrying transcriptional consequence. Conversely, *JAK2*, which showed no differential methylation (Fig. 2), displayed the second-strongest coupling in the panel (cg03508284, ρ = −0.31, padj = 6.5 × 10⁻¹¹), indicating that its lack of differential methylation is unlikely to reflect insensitivity of the assay at this locus.

### Transcriptomic separation from control increases with exposure burden

Hierarchical clustering of differentially expressed transcripts showed progressively greater separation from controls with increasing exposure burden: exposed and control samples were largely intermixed in the single-exposure comparison, a small subset of exposed samples clustered separately in the double-exposure comparison, and triple-exposure and control tumors segregated into two well-defined branches (Fig.5). Differentially expressed miRNAs showed a comparable pattern (Supplementary Figure S1).

**Figure 5.**
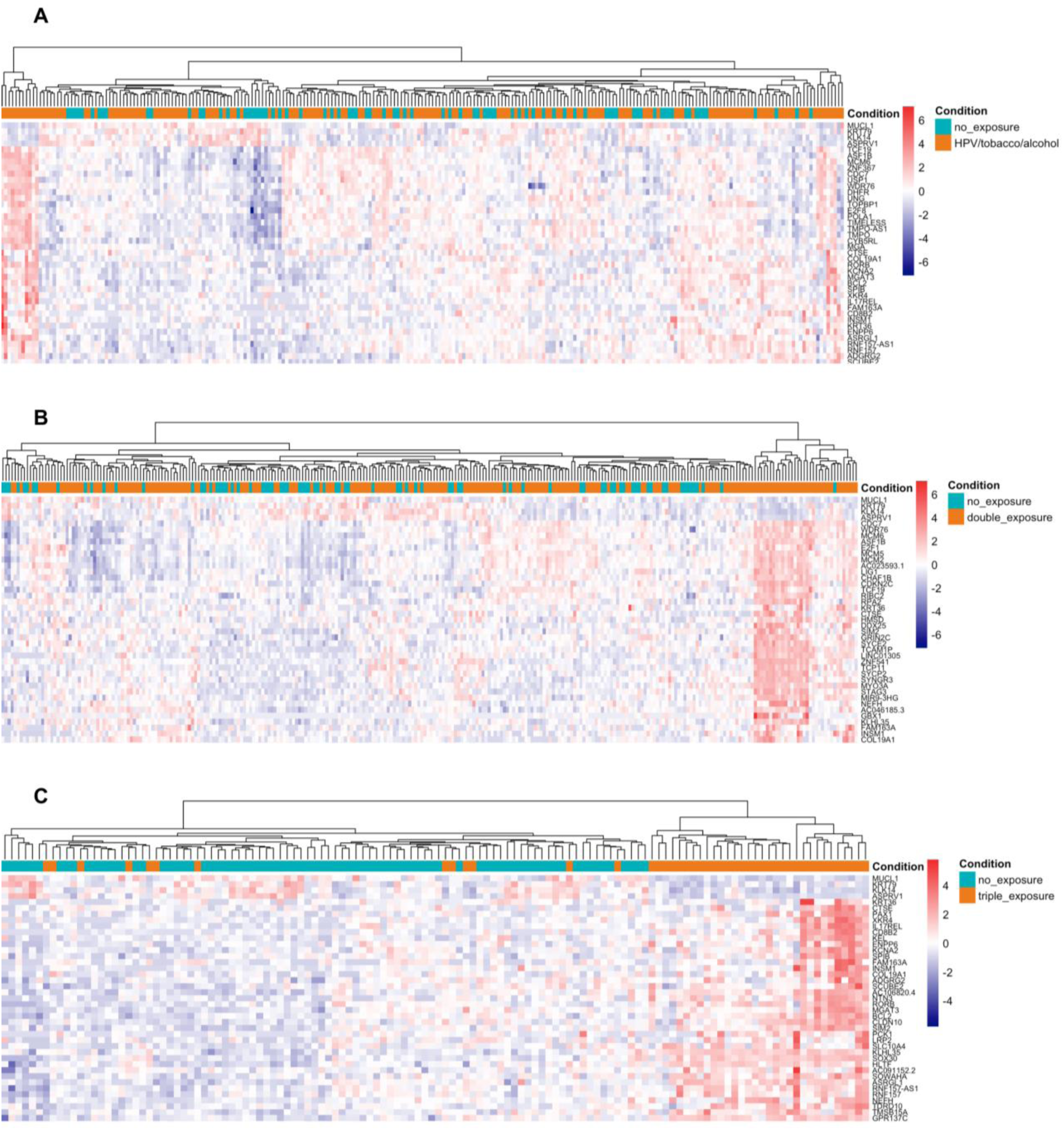
Transcriptomic separation from unexposed controls increases with exposure burden. Hierarchical clustering of differentially expressed transcripts between unexposed control tumors and (**A**) single-exposure, (**B**) double-exposure and (**C**) triple-exposure tumors. Each column represents a patient and each row a transcript. Color indicates row-scaled expression, with red denoting higher and blue lower expression relative to the transcript mean. The condition annotation bar above each heatmap indicates exposure status. Separation between exposed and control samples is minimal in the single-exposure comparison, partial in the double-exposure comparison, and clearly defined in the triple-exposure comparison.

Within this transcriptional landscape, both *CD274* and *JAK2* showed reduced expression in triple-exposure tumors relative to control (*CD274*: 1.51-fold reduction, padj < 0.05; *JAK2*: 1.44-fold reduction, padj < 0.05), the *CD274* change being consistent with its promoter hypermethylation.

### Triple-exposure tumors show increased estimated immune infiltration alongside reduced immune activation signatures

To characterize the immune context of these transcriptional changes, we estimated the relative abundance of tumor-infiltrating immune populations from bulk expression data. Tumors with higher exposure burden, particularly in the double- and triple-exposure subgroups, showed greater estimated infiltration of CD8⁺ T cells, activated memory CD4⁺ T cells and regulatory T cells relative to controls, with the highest estimates in the alcohol and HPV and triple-exposure subgroups (Fig. 6A-C).

**Figure 6.**
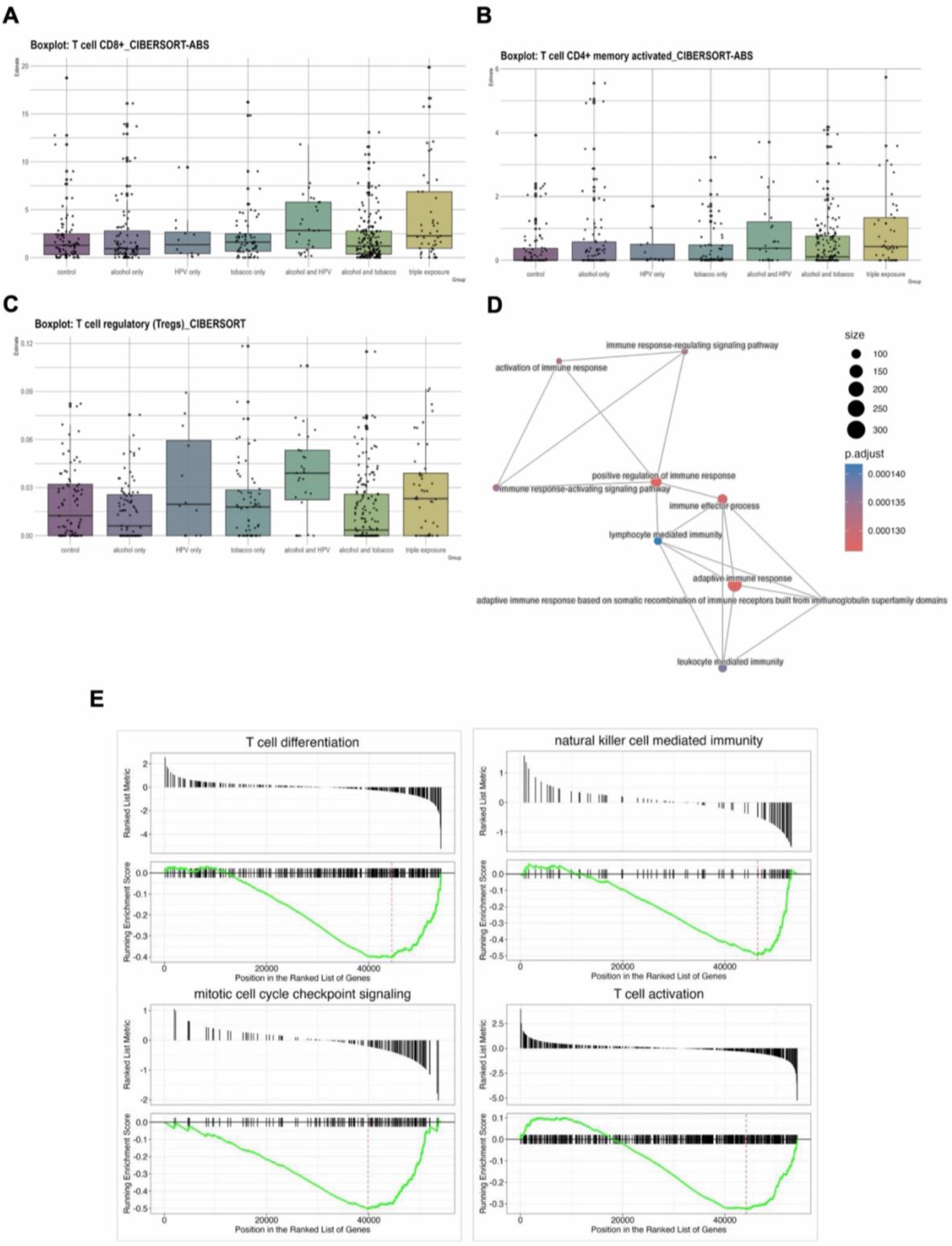
Immune microenvironment of exposure subgroups. (**A**–**C**) Estimated abundance of selected tumor-infiltrating immune populations across exposure groups, inferred from bulk RNA-seq using the TIMER2.0 platform. Each point represents one patient; boxes indicate interquartile range and whiskers the overall distribution. (**D**) Network representation of significantly enriched immune-related Gene Ontology terms. Node size is proportional to gene set size, node color to adjusted p value, and edges connect terms sharing member genes. (**E**) Gene set enrichment analysis of the triple-exposure versus control comparison for representative immune-related Gene Ontology Biological Process terms. Upper subpanels show the ranked list metric across all genes; lower subpanels show the running enrichment score, with vertical ticks marking the position of gene set members within the ranked list. Negative enrichment scores indicate coordinate downregulation in triple-exposure tumors.

In contrast, gene set enrichment analysis of the triple-exposure versus control comparison showed coordinate downregulation of immune activation programs, including T cell activation, T cell differentiation and natural killer cell-mediated immunity, as well as mitotic cell cycle checkpoint signaling (Fig. 6E). The downregulated gene sets were interconnected and spanned adaptive and innate immunity, including adaptive immune response, immune effector process, lymphocyte- and leukocyte-mediated immunity, and immune response-activating signaling (Fig. 6D; all padj < 0.001).

Together, these analyses suggest that tumors with higher exposure burden harbor more immune cells but reduced expression of the programs through which those cells exert effector function, consistent with an inflamed but functionally suppressed microenvironment.

### Somatic mutation and copy number alteration in the checkpoint axis

Because loss-of-function mutations in JAK–STAT components are an established mechanism of checkpoint blockade resistance [11,12], we asked whether somatic alteration could explain the differences in triple-exposure tumors. Mutation frequencies across the interferon-γ–PD-L1 axis were low, and none differed significantly between groups (Figure 7A–B). JAK1 was mutated in two of 82 controls and no triple-exposure tumors, STAT1 in two triple-exposure tumors and no controls, and no patient carried a CD274 mutation. PIK3CA was the most frequently mutated gene in both groups (12 of 45 triple-exposure tumors vs. 11 of 82 controls), again without significant difference.

**Figure 7.**
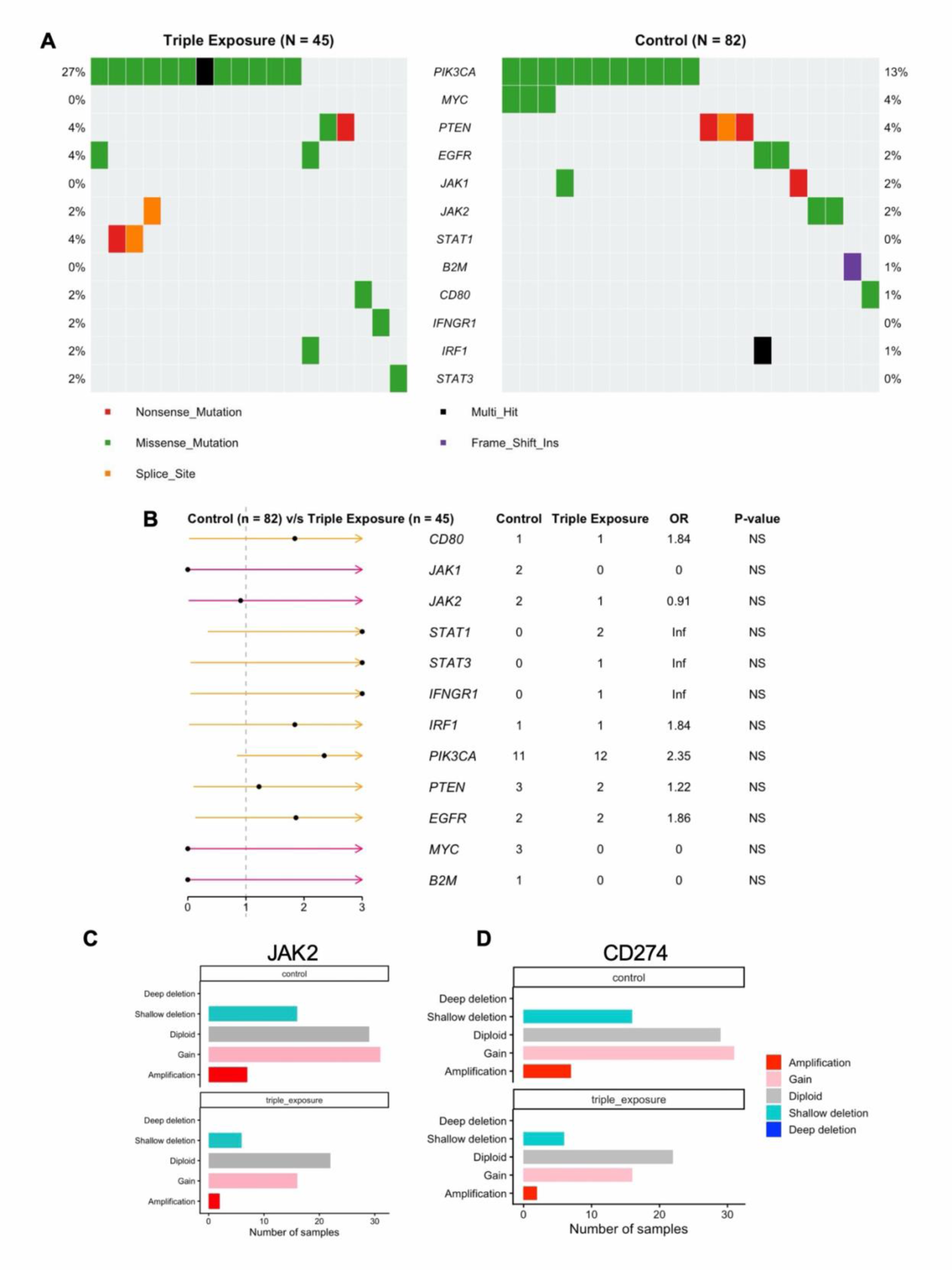
Somatic mutation and copy number alteration in the interferon-γ–PD-L1 axis. (**A**) Co-oncoplot of mutations in checkpoint and interferon pathway genes in triple-exposure (n = 45) and unexposed control (n = 82) tumors. Each column represents a patient and each row a gene; color indicates variant class, and percentages give the proportion of mutated samples in each group. (**B**) Odds ratios with 95% confidence intervals for the comparison of mutation frequency between triple-exposure and control tumors. Values < 1 indicate a higher frequency in control tumors; Inf indicate mutations are only present in the triple exposure group. Mutation counts per group are shown alongside each gene. (**C**) *JAK2* copy number states in control and triple-exposure tumors, expressed as the number of samples in each category. (**D**) *CD274* copy number states in control and triple-exposure tumors.

Copy number states of CD274 and JAK2 were comparable between groups, with most tumors diploid or gained and no significant difference for either gene (padj = 0.42), so reduced CD274 expression does not appear attributable to copy number loss.

## Discussion

In this study, we examined promoter methylation across seven PD-L1-related genes in HNSCC tumors stratified by every observed combination of alcohol, tobacco and HPV exposure. Significant differential methylation was confined to subgroups in which HPV was present and was most extensive in tumors carrying all three exposures, in which six of the seven candidate genes were affected. Among these, *CD274* showed the clearest link between methylation and transcription, with promoter hypermethylation in triple-exposure tumors and an inverse relationship between methylation and expression across the cohort. Triple-exposure tumors additionally showed greater estimated immune infiltration alongside reduced expression of immune activation programs, and neither somatic mutation nor copy number alteration at the loci examined offered a genetic explanation for these differences.

### The role of HPV

The restriction of significant findings to HPV-containing subgroups is the most consequential feature of these results, although its interpretation remains uncertain. HPV-positive HNSCC is known to exhibit a distinct, broadly hypermethylated genome relative to HPV-negative disease [5,15], which may partly explain these findings. Notably, PDCD1 promoter methylation has itself been associated with HPV status in HNSCC [18]. However, the observed changes were directionally heterogeneous rather than uniformly hypermethylating: CD274, IRF1, JAK1 and MIR34A gained methylation, while PDCD1 and MIR33A lost it. This pattern is inconsistent with a purely global hypermethylation phenotype. Moreover, the number of significantly altered loci and breadth of change increased from HPV-only tumors to alcohol–HPV and triple-exposure tumors, suggesting behavioral exposures may modify the extent of an HPV-associated methylation phenotype.

The present analysis compared each exposure group with unexposed controls, and a direct contrast between triple-exposure and HPV-only tumors falls outside its scope. With thirteen patients in the HPV-only subgroup, such a comparison would in any case be limited in power. The interpretation these data best supports is therefore that HPV infection appears necessary for the methylation phenotype described here, and that concurrent alcohol and tobacco exposure are associated with its extent. Fully separating modification from co-occurrence will benefit from cohorts in which HPV-positive tumors with and without behavioral exposure are represented in comparable numbers.

### CD274 methylation and transcript abundance

Among the genes examined, *CD274* offers the clearest link between methylation and transcriptional output. Promoter methylation at cg19724470 was inversely correlated with *CD274* expression across the full cohort (ρ = −0.48), and the same probe was the most significantly hypermethylated *CD274* site in triple-exposure tumors. Reduced *CD274* expression in this group is therefore concordant with the direction of the methylation change and with the cohort-wide relationship between the two measures. Promoter methylation has previously been reported to modulate PD-L1 expression in several malignancies [16,17], and our findings are consistent with that literature.

Three features of the analysis define how this observation is best read. The specificity of the finding is supported by the behavior of the other panel members: *IRF1* and *JAK1* showed significant, albeit small, methylation changes at probes that were uncorrelated with their own expression, meaning that *CD274* is the only locus in the panel at which exposure-associated methylation change and transcriptional coupling coincide. The correlation was computed across the full cohort rather than within exposure groups, and it therefore establishes that the locus behaves as a methylation-responsive promoter in this tissue, which provides a mechanistic context for the group difference rather than a direct test of it. Additionally, *CD274* was supported at the probe level, but did not appear among the significant differentially methylated regions, so the evidence for this locus rests on single-CpG resolution. Region-level replication in an independent cohort would be a natural next step.

### Methylation at PDCD1

*PDCD1* showed promoter hypomethylation at the probe level and gene-body hypermethylation at the region level in the same contrast. Because regions were called genome-wide without restriction to promoter intervals and subsequently assigned to genes, these results describe distinct genomic compartments within the same locus, and discordance between promoter and gene-body methylation has been described previously [18,41]. We therefore report both rather than reconciling them, and do not interpret *PDCD1* as showing a single directional change.

An additional consideration applies to this gene in particular. *PDCD1* is expressed predominantly by infiltrating lymphocytes rather than by tumor cells, and bulk tumor DNA accordingly reflects a mixture of tumor and immune cell genomes whose proportions vary between samples [42]. Given that greater estimated immune infiltration was observed in the same subgroups in which *PDCD1* appears hypomethylated, and that lymphocytes carry a demethylated PDCD1 promoter [43], cellular composition may contribute to this signal. The comparatively weak methylation–expression correlation observed for *PDCD1* (ρ = −0.11) is consistent with that possibility. This locus is therefore interpreted more conservatively than *CD274*, and its behavior would be clarified by analyses incorporating tumor purity or cell-type deconvolution of the methylation data.

Notably, this consideration does not apply in the same way to *CD274*. Triple-exposure tumors showed greater estimated immune infiltration alongside increased rather than decreased *CD274* promoter methylation, so a compositional account would predict the opposite of what was observed. The *CD274* finding is correspondingly more robust to this particular influence.

### An inflamed but functionally suppressed microenvironment

Immune deconvolution and gene set enrichment analysis produced results that are superficially opposed: tumors with higher exposure burden showed greater estimated infiltration of several immune populations while simultaneously showing coordinate downregulation of immune activation programs, including T cell activation, T cell differentiation and natural killer cell-mediated immunity. Taken together, these observations describe a microenvironment in which immune cells are present but the transcriptional programs through which they exert effector function are attenuated.

This configuration is relevant to checkpoint blockade because it dissociates two features that PD-L1-based biomarkers implicitly assume travel together. Tumor PD-L1 expression is physiologically induced by interferon-γ released from activated lymphocytes, so PD-L1 positivity is generally interpreted as evidence of an active, interferon-exposed microenvironment [9,44,45]. Our findings suggest that in tumors carrying combined exposure, the presence of infiltrate need not indicate an intact interferon response, and reduced PD-L1 expression need not indicate an immunologically inert tumor. Exposure history may therefore be a relevant variable in interpreting checkpoint status, although establishing whether it predicts treatment response will require cohorts with recorded immunotherapy outcomes.

### Absence of a genetic explanation

Loss-of-function mutation in JAK–STAT pathway components is an established mechanism of acquired resistance to checkpoint blockade [11,12], and represents the leading alternative explanation for the pathway differences described here. We found no evidence for it. Mutation frequencies at these loci were low in both groups and none differed significantly, with *JAK1* mutated only in control tumors and *STAT1* only in triple-exposure tumors, in both cases at counts too small to interpret. *CD274* copy number states were likewise comparable between groups, with no deep deletions in either, indicating that reduced *CD274* expression in triple-exposure tumors is unlikely to reflect genomic loss at the locus. While these analyses do not themselves demonstrate an epigenetic mechanism, they are consistent with one, and they address the most direct genetic alternative.

### Limitations

Several considerations define the scope of these findings. The study is cross-sectional and observational, and the relationships described are associative rather than causal. Exposure was classified dichotomously from clinical records, which capture the presence of alcohol and tobacco history but not intensity, duration or recency; dose–response relationships therefore remain to be examined, and some degree of exposure misclassification is possible. HPV status was drawn from clinical annotation, and cohorts with uniformly applied molecular HPV testing would allow this classification to be refined. Two of the three HPV-containing subgroups were modest in size (n = 13 and n = 28), and no patient in this cohort carried tobacco and HPV exposure without alcohol, so that particular combination could not be evaluated.

The exposure groups also differ in tumor characteristics beyond the exposures themselves. HPV-positive HNSCC differs systematically from HPV-negative disease in anatomic site, stage at presentation and patient age, and these factors may contribute to the differences reported here; models incorporating them would help to isolate the contribution of exposure specifically. Tumor purity and cellular composition were likewise not incorporated into the present models, and as discussed above in relation to *PDCD1*, bulk methylation at immune-expressed loci is sensitive to the proportion of infiltrating cells in a sample. Immune abundances were inferred computationally rather than measured directly, and deconvolution estimates carry corresponding uncertainty. Significance for the candidate panel was assessed by correction within 51 pre-specified probes, which is appropriate for a targeted analysis while being less stringent than a genome-wide threshold; genome-wide results are reported separately. Although matched miRNA sequencing data were available, promoter methylation at *MIR33A* and *MIR34A* was not examined in relation to the abundance of mature miR-33a and miR-34a, and the transcriptional consequences of methylation at these loci therefore remain to be established. Finally, replication in an independent methylation cohort, assessment of PD-L1 at the protein level, and analysis in immunotherapy-treated patients each represent natural extensions of this work, and inferences regarding checkpoint blockade are at present indirect.

## Conclusion

Combined alcohol, tobacco and HPV exposure is associated with coordinated promoter methylation across the interferon-γ–PD-L1 signaling axis in HNSCC, with the clearest evidence at *CD274*, where promoter hypermethylation corresponds to reduced transcript abundance. These changes were confined to tumors in which HPV was among the contributing exposures and were accompanied by an immune microenvironment showing infiltration without corresponding activation. Neither somatic mutation nor copy number alteration at the loci examined provided a genetic explanation for these differences. Together these observations identify DNA methylation as a candidate mechanism linking exposure history to checkpoint pathway regulation, and suggest that how a patient’s tumor arose may be relevant to how its checkpoint status should be read.

## Ethics statement

This study used publicly available, de-identified data generated by The Cancer Genome Atlas, for which informed consent was obtained by the contributing institutions. Controlled-access sequencing data were obtained through dbGaP (accession phs000178) under an approved data access request. No additional ethical approval was required for this secondary analysis [confirm with your IRB/institutional policy].

## Data availability statement

DNA methylation, clinical and somatic mutation data for the TCGA-HNSC cohort are publicly available from the Genomic Data Commons (https://portal.gdc.cancer.gov/). Raw mRNA and miRNA sequencing reads are available through dbGaP under accession phs000178. Copy number data are available from cBioPortal (https://www.cbioportal.org/). Immune deconvolution estimates were obtained from TIMER2.0 (http://timer.cistrome.org/).

## Code availability

The code used for the analyses in this study is available from the corresponding author upon reasonable request

## Supplementary materials

Supplementary Figure S1 is provided with the current preprint. Additional supplementary figures and tables are in preparation and will be added in a subsequent version.

