## Supplementary materials for "Alcohol, Tobacco, and HPV exposure synergistically regulate promoter methylation of PD-L1 related genes in Head and Neck Squamous Cell Carcinoma"

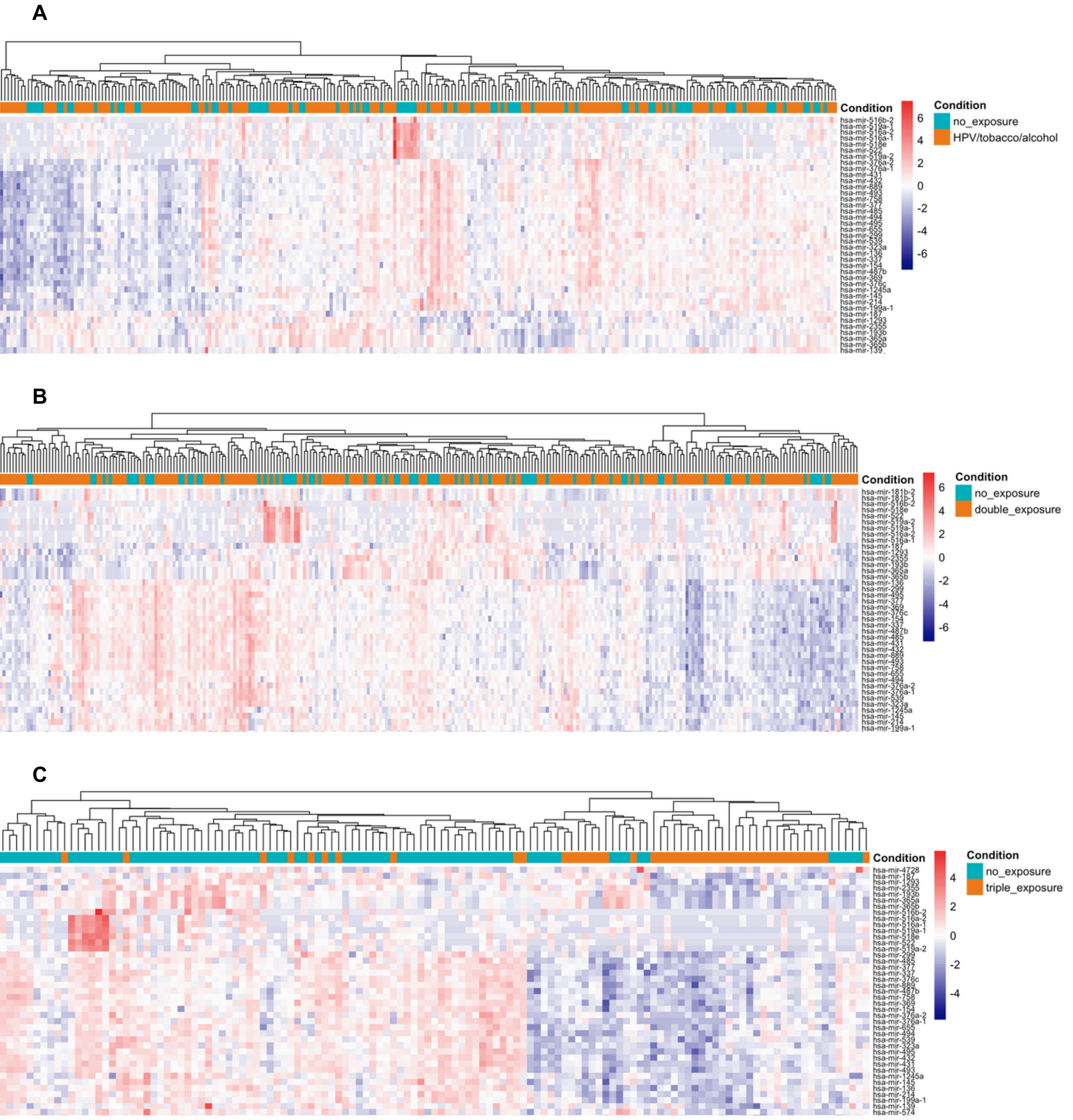


**Supplementary Figure S1. Differential miRNA expression across exposure subgroups.** Hierarchical clustering of differentially expressed miRNAs between unexposed control tumors and (A) single-exposure, (B) double-exposure and (C) triple-exposure tumors. Color indicates row-scaled expression.
